# FishSCT: a zebrafish-centric database for exploration and visualization of fish single-cell transcriptome

**DOI:** 10.1101/2022.09.21.508858

**Authors:** Cheng Guo, Weidong Ye, You Duan, Wanting Zhang, Yingyin Cheng, Mijuan Shi, Xiao-Qin Xia

## Abstract

With the advancement of single-cell sequencing technology in recent years, an increasing number of researchers have turned their attention to the study of cell heterogeneity. In this study, we created a fish single-cell transcriptome database centered on zebrafish (*Danio rerio*). FishSCT currently contains single-cell transcriptomic data on zebrafish and 8 other fish species. We used a unified pipeline to analyze 129 datasets from 44 projects from SRA and GEO, resulting in 964/26,965 marker/potential marker information for 245 cell types, as well as expression profiles at single-cell resolution. There are 117 zebrafish datasets in total, covering 25 different types of tissues/organs at 36 different time points during the growth and development stages. This is currently the largest and most comprehensive online resource for zebrafish single-cell transcriptome data, as well as the only database dedicated to the collection of marker gene information of specific cell type and expression profiles at single-cell resolution for a variety of fish. A user-friendly web interface for information browsing, cell type identification, and expression profile visualization has been developed to meet the basic demand in related studies on fish transcriptome at the single-cell resolution.

## Introduction

Researchers can use scRNA-seq technology to conduct transcriptome studies at the single-cell level (Regev et al. 2017), as well as resolve the types and heterogeneity of cells in tissues. Because of the rapid development and popularization of this technology, a large amount of single-cell transcriptome data has been generated. Even though studies on scRNA-seq have focused primarily on mammals like humans and mice, studies on other model species have also gathered a significant amount of transcriptome data at the single-cell level, particularly in zebrafish (*Danio rerio)*. Zebrafish, as a lower vertebrate, has 87% of the human protein sequence (Klee et al. 2012) and is simple to reproduce, feed, and genetically manipulate on a large scale. Because of its many advantages, it has become one of the most studied model species. It is widely used in developmental, disease model, regeneration, cancer, and evolution research (Parichy 2015). scRNA-seq has been widely used to investigate the transcriptional heterogeneity of zebrafish in various tissues at various time points (Raj et al. 2020; Tatarakis et al. 2021), covering a wide range of experimental types, including baseline (Xu et al. 2020), disease (Campbell et al. 2021; Nayar et al. 2021), regeneration (Navajas Acedo et al. 2019), and so on. scRNA-seq enables large-scale studies of gene expression patterns or marker genes in specific cell types. The rapid growth of data necessitates the development of new databases to accommodate and mine meaningful information, as well as to make the data freely available to interested researchers. Furthermore, for cell type identification, a critical step in scRNA-seq analysis, researchers frequently need to use one or several genes with specific expression patterns as clues and consult a large number of articles for cell type annotation. This approach presents a significant workload challenge, and the identified cell types are not reliable when evidence is limited or even insufficient. Databases can easily sort out the evidence information related to marker gene in a large number of published scRNA-seq projects, and provide easy access and use of this information, which will undoubtedly and greatly improve the convenience and accuracy of related studies about single-cell transcriptome.

The majority of existing single-cell transcriptome databases focus on human and mouse data, such as SC2disease (Zhao et al. 2021) and CancerSCEM (Zeng et al. 2022), which are associated with human diseases, and scRNASeqDB (Cao et al. 2017), PanglaoDB (Franzen et al. 2019) and scAPAatlas (Yang et al. 2022), which contain data from both humans and mice. There are also databases that contain scRNA-seq data from other species, such as DRscDB for Drosophila (Hu et al. 2021) and PlantscRNAdb for a variety of plants (Chen et al. 2021). However, despite being an important multipurpose model species, researchers do not have access to a zebrafish single-cell omics database. There are only a few zebrafish datasets in several databases, including Expression Atlas (Moreno et al. 2022) and DRscDB (Hu et al. 2021). Worryingly, there is no single-cell transcriptome database for fish, the largest vertebrate group (Britz 2017). As a result, the creation of a single-cell transcriptome database for fish, particularly zebrafish, is not only necessary for biomedical research, but also advantageous for the advancement of fish genetic breeding research.

In this study, data were collected from published articles related to scRNA-seq in zebrafish and a variety of other fishes, and the raw data were downloaded from two public data platforms, GEO and SRA. Through the unified pipeline analysis, we obtained a large number of marker/potential marker information of various cell types, as well as expression profiles at single-cell resolution. In addition, we developed FishSCT, a user-friendly and interactive fish single-cell transcriptome data platform. At the moment, this is the largest and most comprehensive online resource for zebrafish single-cell transcriptome data, as well as the only database dedicated to the collection of marker gene information of specific cell type and expression profiles at single-cell resolution for a variety of fish.

FishSCT provides a simple visual interface for searching for gene expression profiles in specific cell types. Furthermore, FishSCT’s multi-search service allows users to obtain marker and expression information for multiple genes at the same time, making it much easier to identify cell types in scRNA-seq analysis.

## Results

### Database contents

At present, FishSCT contains the information of 964 marker genes and 26,965 potential marker genes for 245 cell types, and a large number of single-cell frequency expression profile data (cell number: 646,641). Among them, zebrafish related data account for the vast majority, including 848 markers and 13,800 potential marker for 222 cell types. The numbers of cell types and markers of each zebrafish tissue/organ are shown in Figure 1.

**Figure 1.**
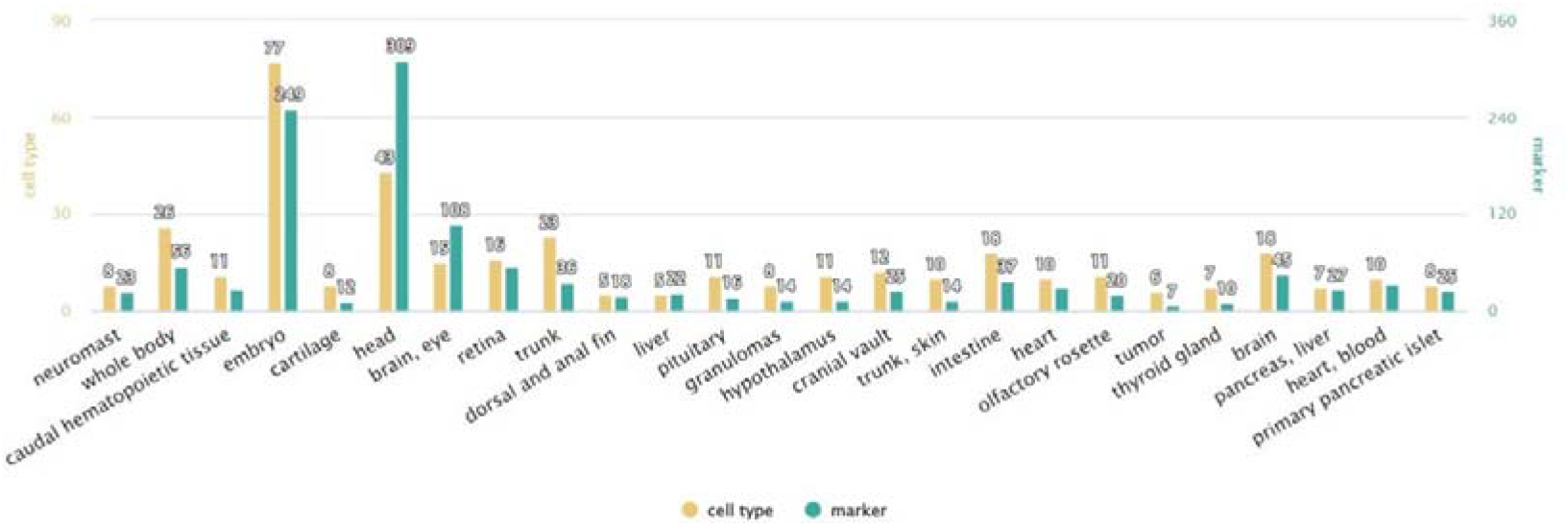
The number of cell types and marker genes in each tissue/organ type of zebrafish.

FishSCT includes single-cell transcriptome information on the model species medaka (*Oryzias latipes*) as well as 7 important economic or ornamental fish species in addition to zebrafish. We obtained 116/13,165 marker/potential marker gene information for 58 cell types of these 8 fish species through unified analysis and mining of public datasets (Figure 2).

**Figure 2.**
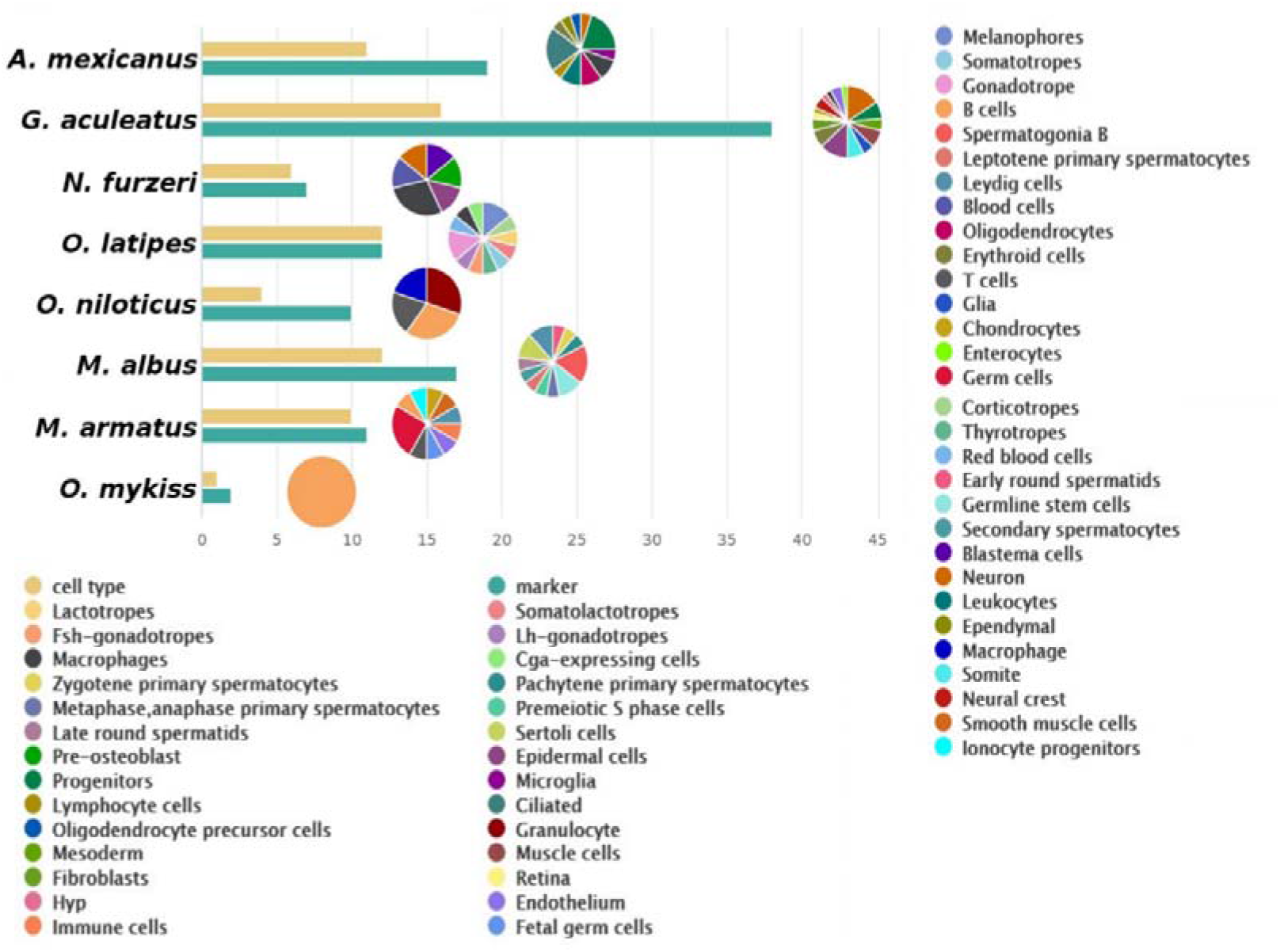
The number of cell types and marker genes in 8 species covered by FishSCT, and the proportion of each cell type.

### FishSCT function and web interface

The six main page sections of FishSCT’s website are “Home”, “Search”, “Multi-search”, “BLAST”, “Statistics”, and “Download” (Supplementary Figure S1). It primarily has the following features in terms of functions: search, visualization, multi-search, statistics and download.

#### Manners of searching

The search function consists primarily of comprehensive search, advanced search, and BLAST. Comprehensive search can be conducted by using the search box on the right of the page header. Users can search by species name, tissue/organ name, cell type, gene ID or symbol, project, or dataset. FishSCT will return a list of all results that match the input information, with links in the list leading to the corresponding object’s details page. The advanced search is based on more accurate gene/cell type/project/dataset information, and more options are provided in the form of the “Search” page under the main menu, and the page will return a list of results that match all of the input content. This allows users to retrieve relevant information more accurately (Figure 3). The BLAST tool has been integrated into the main menu’s “BLAST” page (Supplementary Figure S2). Users can set parameters, paste or upload a sequence file, and FishSCT will align the inputted sequences with transcripts from all species and return a list of matching sequences with descending similarity. BLAST outputs are tabular form with comment lines, and FishSCT offers one-click download of aligned sequences.

**Figure 3.**
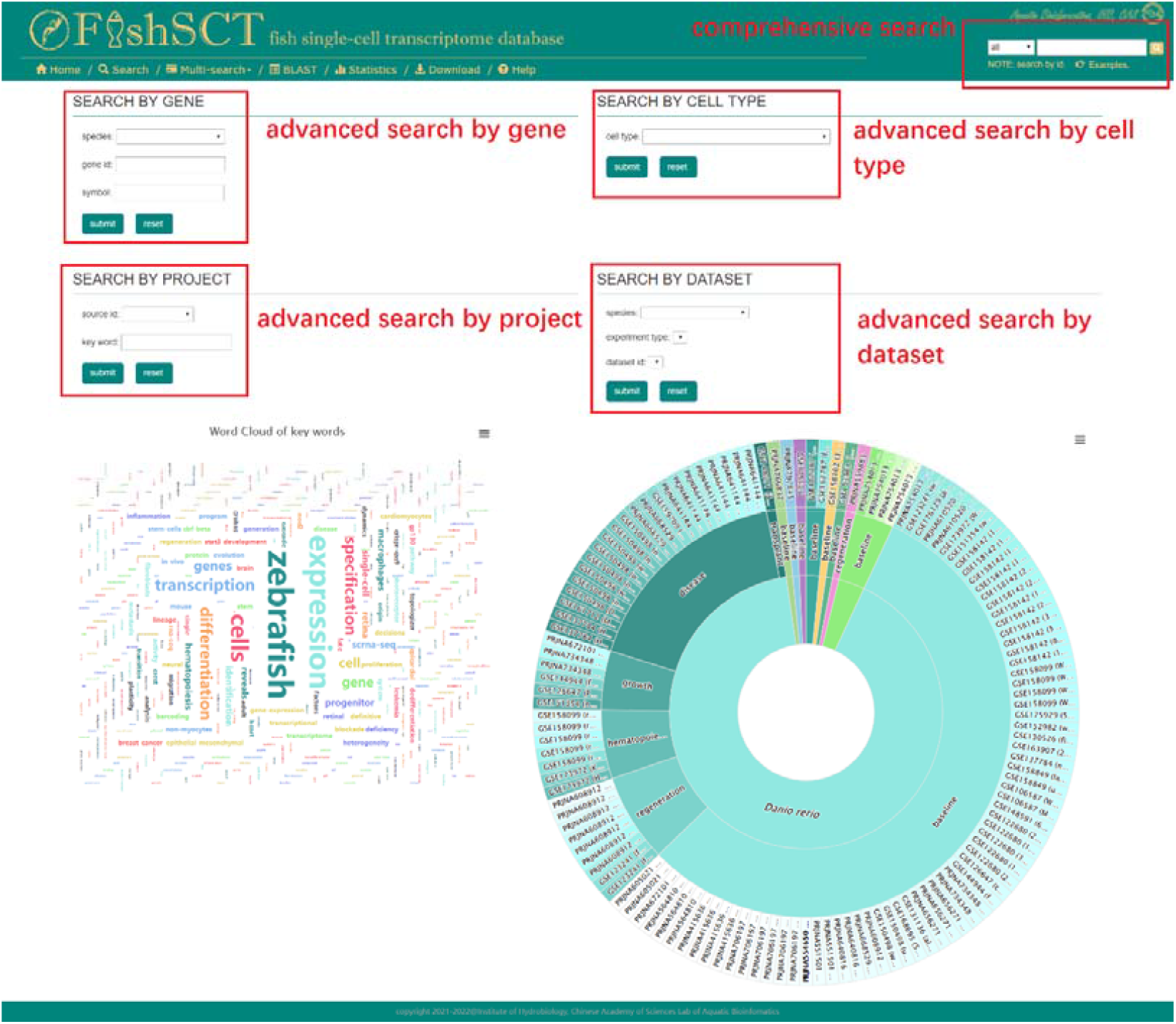
Advanced search page.

#### Presentation of result

The detailed page of a gene, cell type, tissue/organ, species, dataset, or project is returned by the comprehensive or advanced search. The first four pages contain basic information about the source datasets as well as links to related projects. Aside from genes, there are corresponding statistical figures of cell number in the detailed pages of each level to help users understand sample distribution. Furthermore, the word cloud figure is drawn based on the frequency of occurrence of marker gene in the analysis results of each dataset, which can show the cell type’s more statistically significant marker genes.

The gene detail page displays basic information about a gene, such as its symbol, location, gene type, and genome version, as well as a visualization of the structural elements of the gene using the JBrowse2 embedding window. All of the genes queried by users are linked to specific species. If the gene belongs to zebrafish, the page will display a bubble diagram with information about its expression and markers. The diagram’s green bubbles depict the distribution of zebrafish datasets in two dimensions: time point and tissue/organ type. The diamond in the green bubbles indicates that the gene is a marker gene, and the circular dot indicates that the gene is a potential marker gene. To distinguish different cell types, distinct colors are used on both graphic elements. This type of bubble diagram clearly shows which tissues/organs and cell types this gene acts as a marker for (Figure 4A). Furthermore, the magnifying glass button next to each dataset ID can be clicked to view the clustering results of cells in that dataset as well as the expression of this gene (Figure 5).

**Figure 4.**
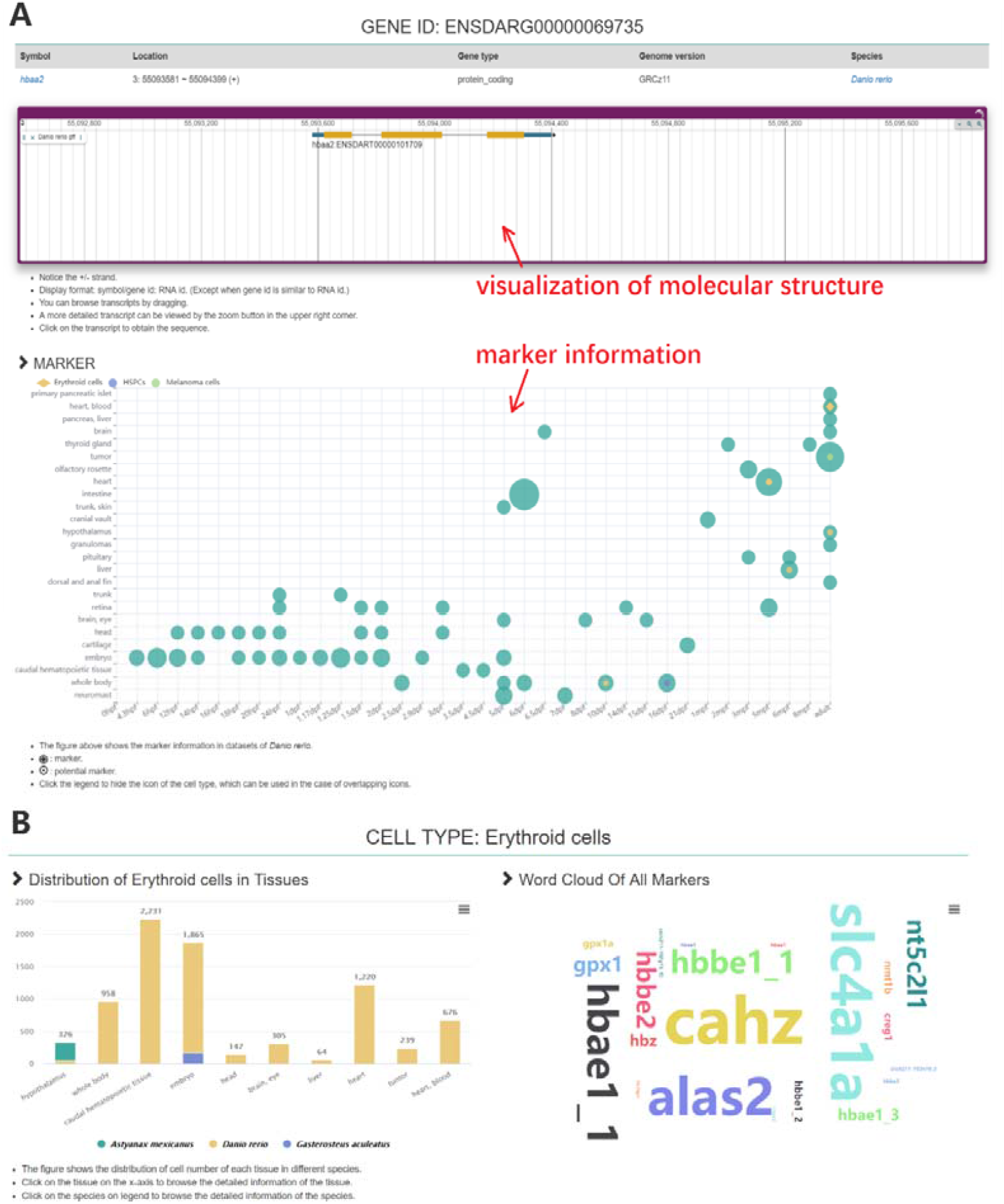
The page for details of gene and cell type. (A) The page for details of gene *hbaa2* (gene id: ENSDARG00000069735). (B) The page for details of cell type erythroid cell.

**Figure 5.**
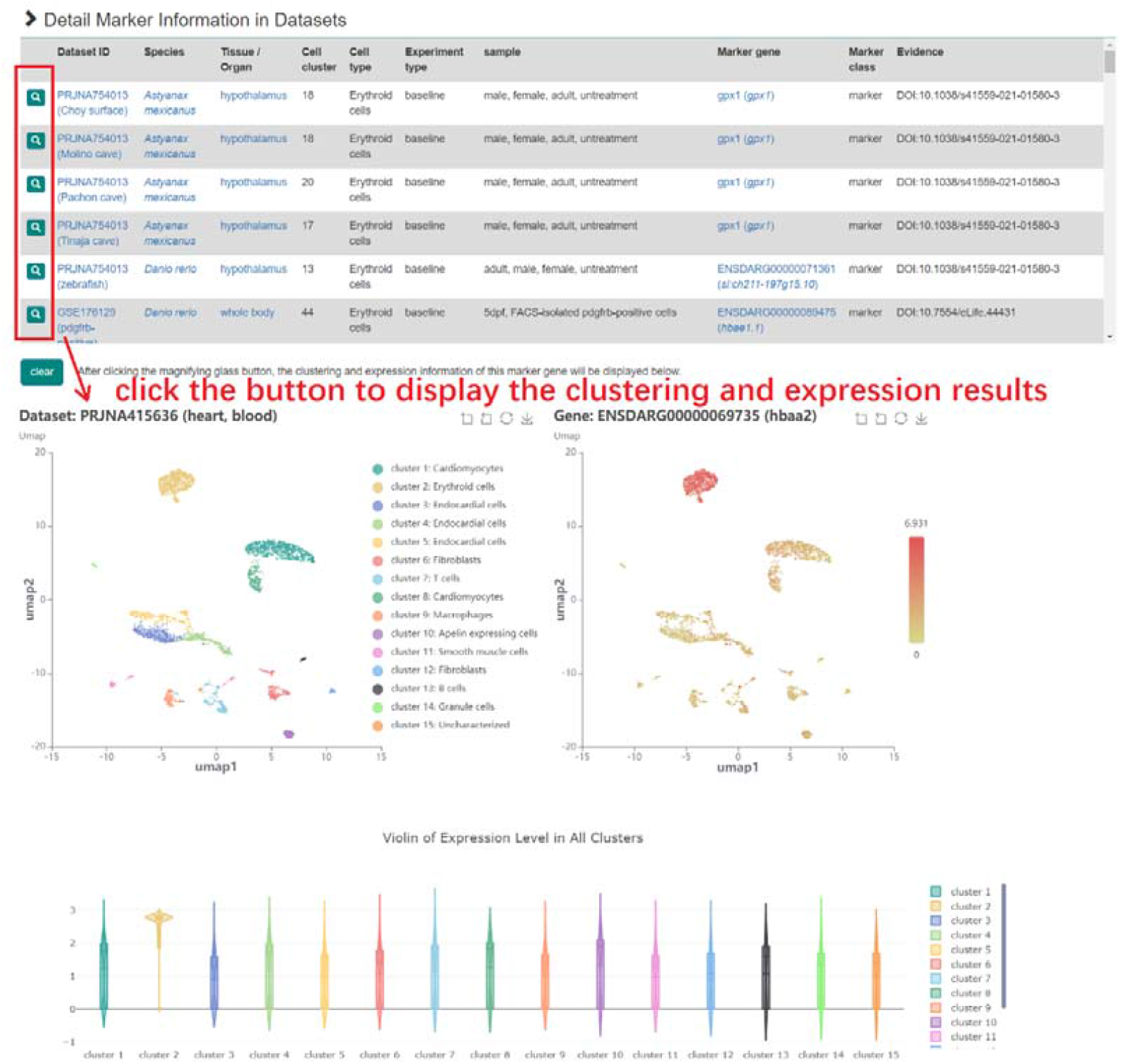
The visualization function of clustering and expression results.

The detailed page of cell type depicts the distribution in various tissues/organs (Figure 4B). More detailed marker gene information for this cell type in each dataset is listed in a table, and users can also browse the cell clustering results of any dataset and the expression of corresponding marker genes, just like on the gene page above (Figure 5). In the tissue/organ detail page, the number of different cell types in this tissue/organ from different species or datasets is displayed in bar charts. The percentages of marker/potential marker genes are presented in a pie chart (Supplementary Figure S3).

In the detailed page of species, the number of different cell types in different tissues/organs is shown as a bar chart. However, research on fish other than zebrafish is limited at this stage, and most studies involve only a single tissue/organ. A pie chart shows the number and proportion of marker/potential marker genes in each cell type. The distribution of dataset numbers for the most data-abundant zebrafish is first shown in two dimensions: time point and tissue/organ type. A bar chart is then used to show the number of cell types and marker genes in each tissue/organ. FishSCT, in addition to listing all datasets for this species, displays the cell number of each type in each dataset via a bar chart (Supplementary Figure S4).

The information about samples and data sources is presented on the dataset’s detail page. Users can not only view the visualization results of cell clustering in this dataset, as well as the number distribution of various cell types in different clusters, but also they can understand the expression of all marker genes in all cell clusters via a bubble chart (Supplementary Figure S5). The project’s detail page includes the article information relevant to this project, as well as a word cloud diagram based on the abstract. This page also contains the clustering results for each dataset as well as the list of marker genes for this project (Supplementary Figure S6).

#### Multi-search function

The two main components of the multi-search function are “Marker” and “Expression”. The former serves as a guide for researchers to identify different cell types, and the latter aids in comparative analysis of expression profiles. Users can enter multiple gene IDs or gene symbols on the “Marker” page under “Multi-Search” in the main menu to get the marker information for these genes (Figure 6A). Users will have a reference for cell type identification thanks to the Sankey diagram’s pointing relationship between cell type, input genes, and marker class (Figure 6B). A table listing the specific restriction information for each marker (gene, species, tissue/organ, cell type, etc.) also includes buttons to display the clustering and gene expression data for the associated datasets (Figure 5). In order to get the desired outcome, users can also set parameters in the form, such as exact or fuzzy matching, p value, etc.

**Figure 6.**
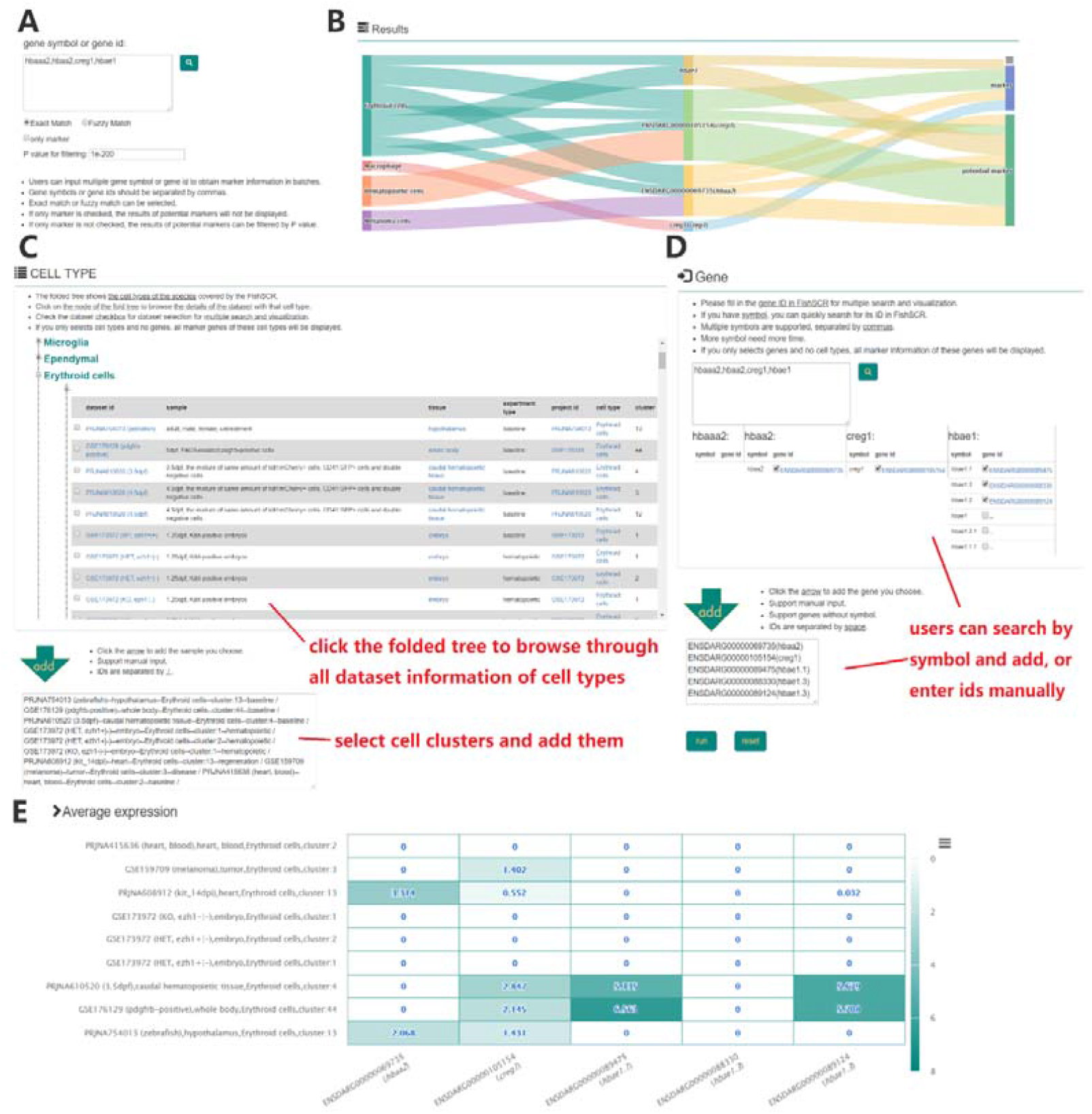
Examples of multi-search function. (A) Input form in “Marker” page. (B) Search results of marker relationship. (C, D) Input form of cell type and gene in “Expression” page. (E) Visualization of expression profile results.

We offer an expression profile visualization service in the “Expression” page under the main menu “Multi-search,” which enables users to select datasets and genes of any species to browse, in order to better display the expression of particular genes in particular cell types. While choosing datasets and genes can be done by manually entering the corresponding IDs, browsing data and making choices during that process is more practical. Users can do this by choosing a target species from the species list on the page’s left and then navigating that species’ cell type through a folded tree in the page’s center. Users can browse the data of the cell cluster of datasets corresponding to the cell type after the nodes of the folded tree’s expanded nodes, including sample description, tissue/organ, experiment type, etc. Through the “add” button underneath the folded tree, users can choose multiple cell clusters from one or more datasets and add them to the list of cell clusters to be visualized (Figure 6C). Users can query and select specific gene IDs in the gene input section, and then add them to the list to be visualized (Figure 6D). The page will return a heat map of related gene expression in these datasets for the user to compare and analyze after clicking the “run” button (Figure 6E). If a marker relationship exists between the cell types of the selected cell clusters and genes, the visualization of corresponding marker information and clustering results will be returned (Figure 5, Figure 6B), as shown on the “Marker” page.

#### Statistics and download

We display some fundamental statistical data of FishSCT on the “Statistics” page of the main menu to help users understand the data content of FishSCT more quickly, such as the number statistics of marker and potential marker genes of each cell type of each fish, the distribution of 129 datasets, and the phylogeny and differentiation period of 9 fish species (Supplementary Figure S7). The main menu’s “Download” page provides free downloads of all marker/potential marker gene data for each species (Supplementary Figure S8). The data table includes the gene ID, symbol, chromosome location, scientific name of the species, marker class, tissue/organ, cell type, p value, and evidence.

## Materials and methods

### Data collation and processing

#### Data source

We manually compiled the published articles about scRNA-Seq in fish and obtained the raw data for 129 datasets across 44 projects from two open data platforms, GEO and SRA, to enable the datasets to cover a wider range of time points and tissue/organ types (Table 1, Figure 7a). Zebrafish datasets, which cover 25 tissues/organs at 36 different time points during the growth and development stages, are the highest number of these datasets, at 117. We divided the datasets into 6 different experiment types, including baseline, disease, regeneration, hematopoietic, growth, and transplant, in accordance with the experiment type and sample status of the published articles. The most are found in the baseline, disease, and regeneration datasets, with 83, 22, and 10 each (Supplementary Figure S9).

**Figure 7.**
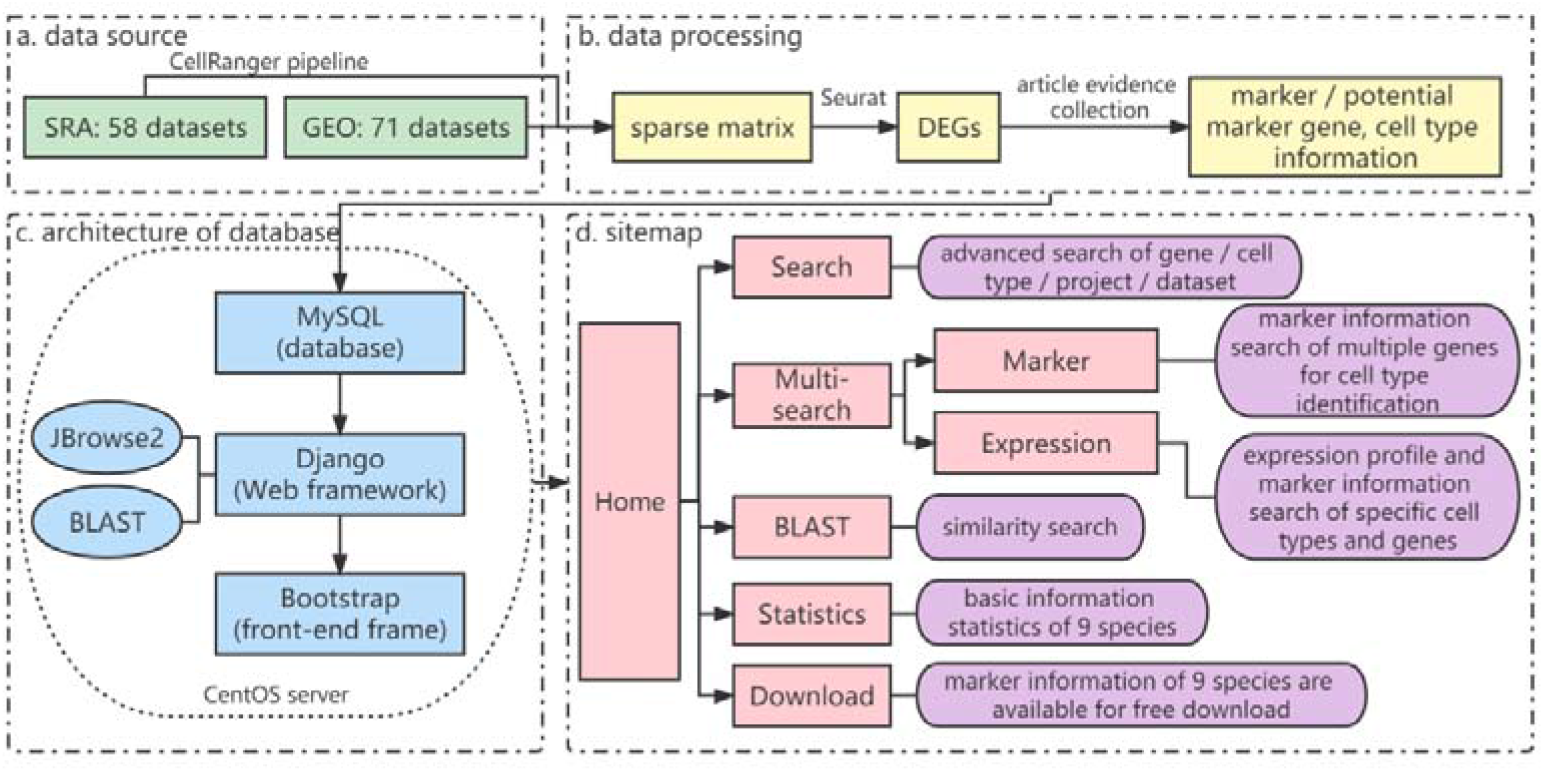
Implementation of FishSCT. (a) Data sources. (b) Data processing. (c) Architecture of FishSCT. (d) Sitemap of FishSCT.

**Table 1.**
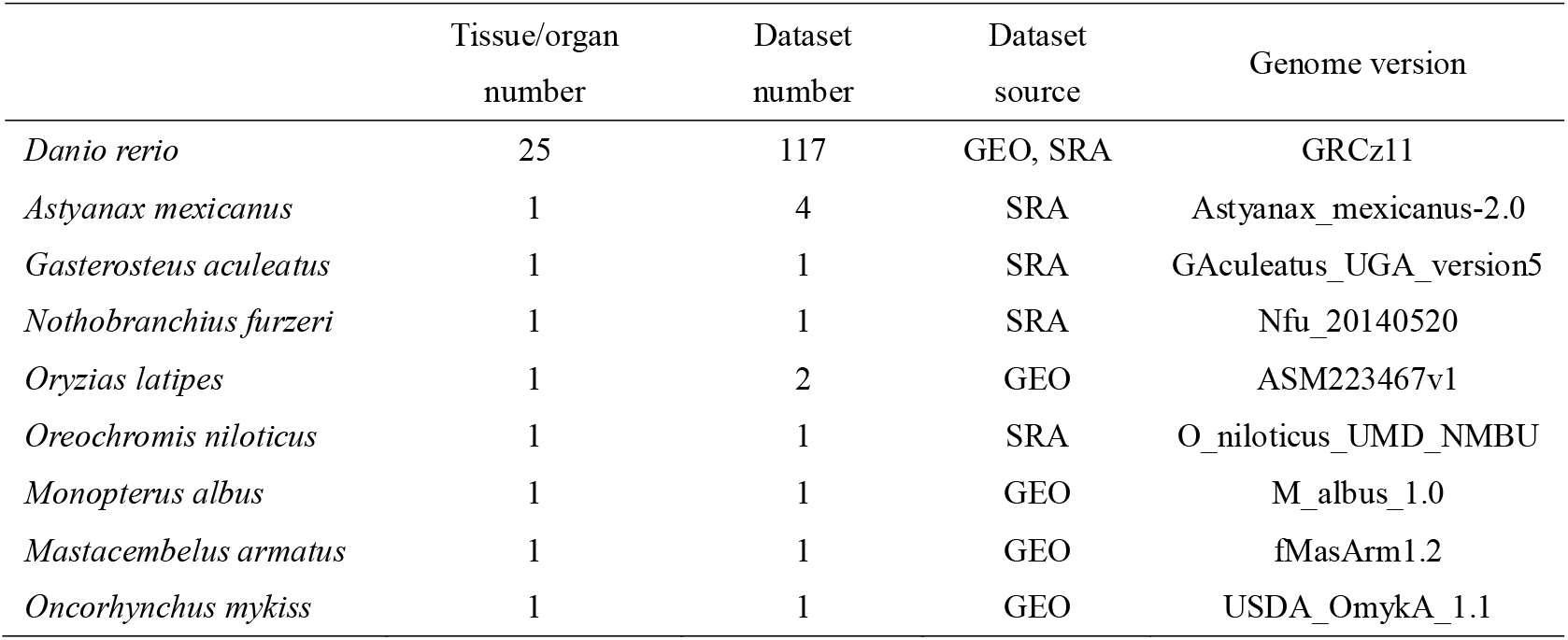
The statistic of datasets

|  | Tissue/organ<br>number | Dataset<br>number | Dataset<br>source | Genome version |
| --- | --- | --- | --- | --- |
| *Danio rerio* | 25 | 117 | GEO, SRA | GRCz11 |
| *Astyanax mexicanus* | 1 | 4 | SRA | Astyanax_mexicanus-2.0 |
| *Gasterosteus aculeatus* | 1 | 1 | SRA | GAculeatus_UGA_version5 |
| *Nothobranchius furzeri* | 1 | 1 | SRA | Nfu_20140520 |
| *Oryzias latipes* | 1 | 2 | GEO | ASM223467v1 |
| *Oreochromis niloticus* | 1 | 1 | SRA | O_niloticus_UMD_NMBU |
| *Monopterus albus* | 1 | 1 | GEO | M_albus_1.0 |
| *Mastacembelus armatus* | 1 | 1 | GEO | fMasArm1.2 |
| *Oncorhynchus mykiss* | 1 | 1 | GEO | USDA_OmykA_1.1 |

#### Data processing

A total of 71 GEO-sourced datasets provide sparse matrices of single-cell expression values that can be directly used for subsequent analysis, and 58 SRA-sourced datasets provide only raw sequencing files, most of which were generated using the 10x Genomics platform. For the latter, we first generated FASTQ files from SRR files using fasterq-dump, then ran the 10x Genomics CellRanger Pipeline (6.1.2) to build custom references and obtain single-cell expression sparse numerical matrices. Seurat (4.1.0) was used for further analysis of sparse numerical matrices (Satija et al. 2015), including data integration, filtering, clustering, cluster refinement and identification of differentially expressed genes in each cluster (min.pct = 0.25, logfc.threshold = 0.25). By referring to marker gene in articles, cell type annotation was carried out on each cluster, and uncertain clusters were labeled as “Uncharacterized” (Figure 7b). We also screened for the potential marker genes, which include all DEGs in clusters containing no more than 20 DEGs, and DEGs with P<1E-50 in other larger clusters. The identification of potential marker gene enriches the results of multi-search function and further helps the identification of cell type. In addition, if the codes or parameters for analysis were provided in articles, we used them in our analysis to follow the article authors’ intentions as much as possible.

### Database construction

#### Database architecture

All of the software tools used in FishSCT are open source (Figure 7c). The website was created with the Python programming language and is based on the Django web framework. The website is hosted on a CentOS server, and the data were imported into a MySQL-based relational database. Services for sequence homology alignment and visualization are provided by BLAST (2.6.0) and JBrowse2. The Bootstrap framework, which offers a dynamic page layout based on the resolution of the user’s display, was used to develop the database’s front-end. The highcharts, echarts, amcharts, and plotly packages were used to implement front-end visualization, which generates interactive charts and offers a user-friendly visualization interface.

#### Database function realization

Django’s ORM model was used to implement the text search function, while in-house Python scripts were used to implement the comprehensive search, advanced search of gene/cell type/project/dataset, and multi-search functions. The BLAST software package is used to perform sequence homology searches. JBrowse2 implemented the visualization of results based on genome location and gene structure. In-house Javascript scripts were used to visualize expression profiles and relationships between markers. The data download service was built using Django’s static file system.

## Discussion

Researchers can now study the transcriptome at the single-cell level on a large scale thanks to scRNA-seq technology. The core content of single-cell analysis is the marker gene. However, most fish species, particularly non-model fish, have few online marker gene resources. FishSCT performed a unified and centralized analysis on the publicly available datasets of 9 fish species, allowing users to easily browse the information of interested marker genes and expression profiles in order to conduct further research on the marker genes or comparative gene expression analyses between different species, experimental types, and developmental stages. Zebrafish, as one of the most widely used model species, holds a central position in FishSCT because it has the most comprehensive data in the database. Its multi-tissue/organ datasets, in particular, cover 36 time points, forming a complete and delicate time line that is extremely valuable for scientific research.

In conclusion, we have built a fish single-cell transcriptome database to keep track of and include pertinent scRNA-seq data from zebrafish and a variety of other fish species, offering a user-friendly online information platform for researchers. We will regularly maintain and update FishSCT to make sure that it continues to serve as a crucial public resource for researchers conducting molecular biology research.

## Acknowledgments

We thank the Analysis and Testing Center at Institute of Hydrobiology for technical supports. This computational work in this study was supported by the Wuhan Branch, Supercomputing Center, Chinese Academy of Sciences, China.

## Data Availability

All data used in this study were derived from GEO or SRA. All data generated and analyzed results were uploaded to FishSCT and provided with corresponding download options. The process and script files can be got through reasonable requests.

## Author Contributions

Cheng Guo: Conceptualization, methodology, formal analysis, investigation, writing—original draft preparation. Weidong Ye: Conceptualization, methodology, formal analysis, investigation. You Duan: Conceptualization, methodology. Wanting Zhang: data curation. Yingyin Cheng: validation, resources, supervision. Mijuan Shi: Conceptualization, investigation, writing—review and editing. Xiao-Qin Xia: Conceptualization, investigation, writing—review and editing, project administration, funding acquisition.

